# Unsupervised mining of HLA-I peptidomes reveals new binding motifs and substantial false positives in community database

**DOI:** 10.1101/2021.09.19.460988

**Authors:** Chatchapon Sricharoensuk, Tanupat Boonchalermvichien, Phijitra Muanwien, Poorichaya Somparn, Trairak Pisitkun, Sira Sriswasdi

## Abstract

Modern vaccine designs and studies of human leukocyte antigen (HLA)-mediated immune responses rely heavily on the knowledge of HLA allele-specific binding motifs and computational prediction of HLA-peptide binding affinity. Breakthroughs in HLA peptidomics have considerably expanded the databases of natural HLA ligands and enabled detailed characterizations of HLA-peptide binding specificity. However, cautions must be made when analyzing HLA peptidomics data because identified peptides may be contaminants in mass spectrometry or may weakly bind to the HLA molecules. Here, a hybrid *de novo* peptide sequencing approach was applied to large-scale mono-allelic HLA peptidomics datasets to uncover new ligands and refine current knowledge of HLA binding motifs. Up to 12-40% of the peptidomics data were low-binding affinity peptides with an arginine or a lysine at the C-terminus and likely to be tryptic peptide contaminants. Thousands of these peptides have been reported in a community database as legitimate ligands and might be erroneously used for training prediction models. Furthermore, unsupervised clustering of identified ligands revealed additional binding motifs for several HLA class I alleles and effectively isolated outliers that were experimentally confirmed to be false positives. Overall, our findings expanded the knowledge of HLA binding specificity and advocated for more rigorous interpretation of HLA peptidomics data that will ensure the high validity of community HLA ligandome databases.

## Introduction

Human leukocyte antigen (HLA) is a family of proteins in the immune system that binds to and presents peptide fragments of proteins expressed in the body for recognition by T cells. Peptides that form stable complexes with HLA proteins are also called HLA ligands. When a foreign antigen, whose amino acid sequence differs from the host’s proteome, was intracellularly processed and presented on the cell surface by HLA proteins, the cell containing foreign antigen would be recognized T cell and subsequently destroyed by the immune system. Therefore, HLA-peptide binding activity has been extensively studied for medical and biotechnology applications in vaccine design and cancer immunotherapy^1-6^.

HLA class I is a subclass of the HLA system that recognizes peptides with 8-15 amino acids in length. The binding affinity of a peptide to an HLA class I molecule mainly depends on an 8-to 10-residue motif on the peptide including a few HLA allele-specific amino acid residues at anchor positions^7-10^. Other residues on the peptide are relatively unconstrained, but some amino acid combinations can affect the binding affinity. To date, although a few works have highlighted the multiple specificities of HLA class I binding^8,11,12^ and HLA class II binding^13^, the motif of each HLA class I allele is still represented with a single amino acid frequency profile in major databases^14,15^. In other words, HLA class I motifs were assumed to be unimodal. While this simplification may not have a noticeable impact on the development of HLA binding prediction models^11,16^, it may limit the design landscape of vaccines if researchers use only the consensus motif as a guideline.

Breakthroughs in HLA peptidomics, which enabled the isolation of HLA proteins from the cell surface followed by high-throughput sequencing of HLA ligands, have cataloged a large amount of ligand sequences for a multitude of HLA class I and class II alleles from both cell lines and patient samples^8,10,17,18^. These data accelerated the improvement in HLA binding prediction accuracy as well as enabled detailed characterization of HLA binding specificity. HLA peptidomics is also being increasingly utilized to identify tumor-specific or tumor-elevated antigens in cancer patients, which can then be developed into a cancer vaccine to boost the immune system to target cancer cells^5,6^. Nonetheless, results from HLA peptidomics only indicate whether the peptides are bound to the HLA proteins and presented on the cell surface but provides no information on their actual binding affinities. Hence, downstream analyses of HLA peptidomics often involve HLA binding affinity predictions by artificial neural network models to screen for peptides with strong bindings. Furthermore, like most mass spectrometry analyses, results from HLA peptidomics can include contaminants such as carry-over peptides and non-HLA-specific proteolytic peptides or artifacts from in-source fragmentations ^19,20^. A recent study has proposed additional analysis steps that would help reduce the number of contaminant identifications originating from these sources^20^.

Increasing the understanding of HLA binding specificity and the quality of known HLA ligand databases is crucial for designing better vaccines against constantly emerging pathogens and improving the accuracy of HLA binding and immunogenicity predictions. In this study, a hybrid *de novo* peptide sequencing strategy with SMSNet^21^ was applied to large-scale HLA class I peptidomics datasets^8,17^ to uncover new candidate HLA ligands that would expand the existing databases. Subsequent unsupervised clustering of known and newly discovered ligands for each HLA class I allele strongly suggested that several alleles recognize multiple, clearly distinct motifs. Many potential false positives whose sequences do not match the corresponding HLA binding motifs were also observed. A validation experiment confirmed that almost all potential false positives exhibit no HLA binding activity. Most importantly, many of these false positives were also found in the Immune Epitope Database^15^ and could be erroneously used by the community. Additionally, our HLA peptidomics analysis of a B-lymphoblastoid cell line expressing both HLA class I and class II alleles highlighted the capability of *de novo* sequencing by SMSNet to identify high-affinity antigens in a multi-allelic setting.

Overall, our work revisited two key aspects of the HLA study: the representation of the HLA binding motif and the interpretation of HLA peptidomics data. The findings strongly suggested that the implicit unimodal assumption of HLA class I motifs should be replaced by a multimodal representation and that the quality of HLA peptidome-derived HLA-I ligands reported in the community database may be questioned.

## Results

### Re-analysis of large-scale mono-allelic HLA class I peptidomes

*De novo* peptide sequencing with SMSNet^21^ was shown to be effective for discovering new candidate HLA class I antigens from a peptidomics dataset. Here, SMSNet was applied to a larger collection of high-quality HLA peptidomics data from mono-allelic human B lymphoblastoid cell lines encompassing 88 HLA-A, -B, -C, and -G alleles^8,17^. In total, 109,372 unique peptide sequences with lengths ranging from 8 to 15 amino acids were identified from 327,312 mass spectra (Figure 1a, Supplementary Table 1). There are 36,043 newly discovered peptide-HLA pairs involving 25,718 unique peptide sequences as well as 5,347 additional pairs that have been previously observed in multi-allelic patient samples. Over 88% (22,854 peptides) of newly discovered peptides could be mapped to the human reference proteome. About half of peptides with unknown origins could be traced to open reading frames on non-coding transcripts (1,630 peptides) and a small fraction could be explained by proteasome-mediated splicing (222 peptides). However, it should be noted that 30% of hypothetical spliced peptides could also be alternatively explained by missense mutations and 45% of them might be erroneously attributed to splicing events (see Methods). The length distribution of newly identified peptides matches well with past observations^22^, with the majority being 9-mers (Figure 1b). Most importantly, the discovery of these new peptides has the potential to expand the database of known HLA class I ligands by up to 35-40% for some major alleles such as HLA-A*11:02 and HLA-A*34:02 (Figure 1c).

**Figure 1.**
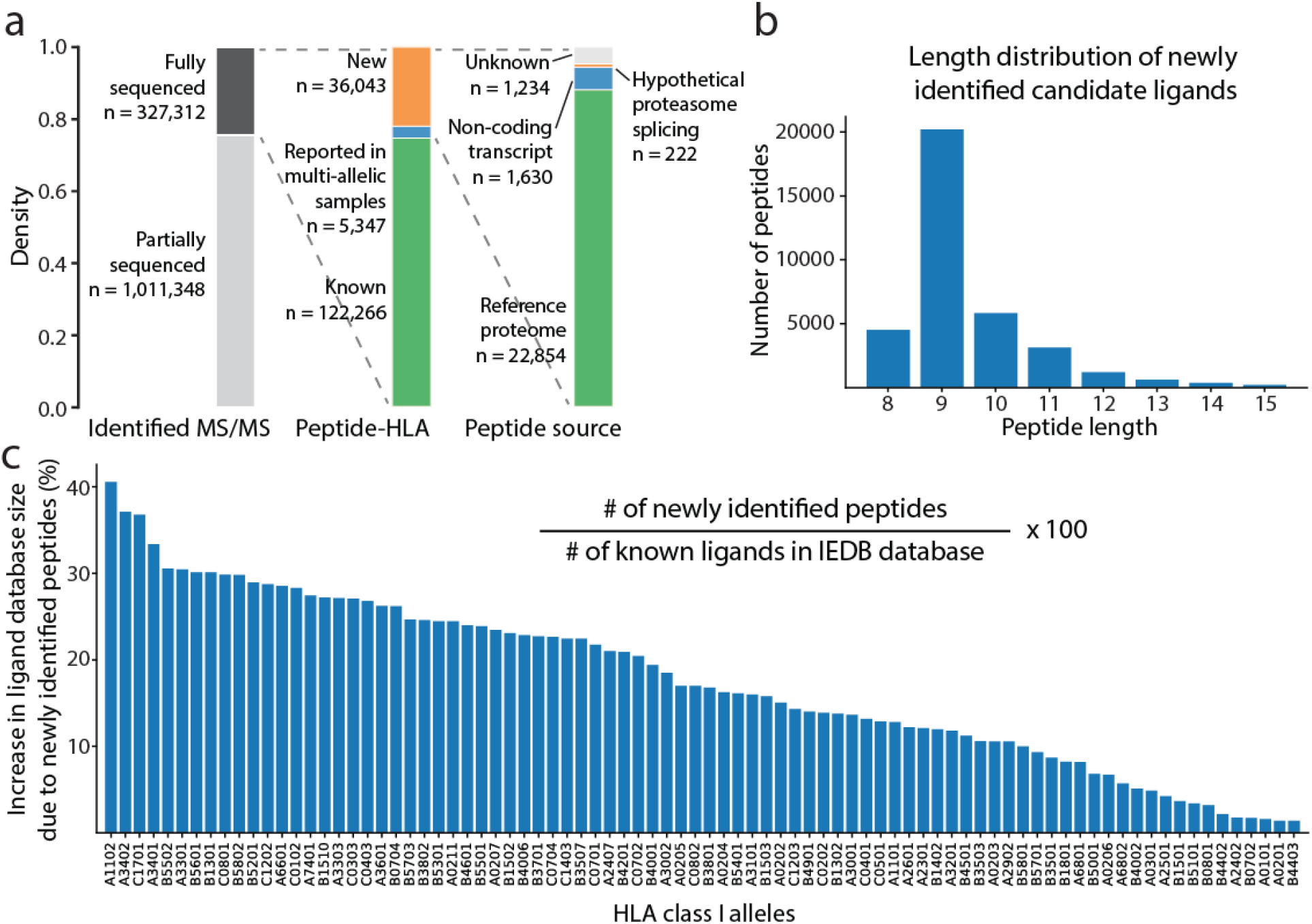
SMSNet identified a large number of new ligands from public HLA peptidomics datasets. a) Statistics of MS/MS spectra, peptide-HLA pairs, and the sources of peptides identified by SMSNet on mono-allelic HLA peptidomics datasets of 88 HLA class I alleles (see Methods). b) Length distribution of all identified peptides. c) Potential increase in the size of the database of known ligands from this study, assuming that all newly identified sequences are true ligands. The number of known ligands for each allele was extracted from the IEDB database by counting unmodified antigens and antigens with major modifications, namely oxidized methionine and phosphorylated serine, threonine, and tyrosine.

### Extent of tryptic peptide contaminations in HLA peptidomics data

Past analyses of HLA peptidomics were careful not to report 9-mer tryptic peptides as antigens for HLA alleles whose binding motifs do not end with an arginine or a lysine^10^. Among 88 HLA class I alleles investigated in this study, 12 have binding motifs ending with an arginine or a lysine (Figure 2a, HLA-A*03:01, HLA-A*11:01, HLA-A*11:02, HLA-A*30:01, HLA-A*31:01, HLA-A*33:01, HLA-A*33:03, HLA-A*34:01, HLA-A*34:02, HLA-A*66:01, HLA-A*68:01, and HLA-A*74:01). However, 2,838 tryptic peptides identified for the other 76 alleles are reported as positive antigens in the Immune Epitope Database (IEDB)^15^. Motif clustering with GibbsCluster^23^ and binding affinity prediction with NetMHCpan^24^ clearly illustrated that these tryptic peptides form a separate cluster with lower binding affinities than the known motifs (Figure 2b and Supplementary Figure 1). Clusters of tryptic peptides were observed for 11 HLA class I alleles where greater than 13% of identified peptides are tryptic. In extreme cases such as for HLA-B*57:01 and HLA-B*35:01, more than 42% of all identified peptides are tryptic, and more than half (365 out of 709) of these tryptic peptides are reported as legitimate ligands in IEDB. To test whether these tryptic peptides are specifically recognized by the corresponding HLA alleles, and thus may be true ligands, predicted binding affinities for observed tryptic peptide-HLA allele pairs were compared with the predicted binding affinities between random pairs. This finding revealed that almost every HLA allele does not exhibit a stronger affinity toward the observed tryptic peptides compared with random tryptic peptides (Supplementary Figure 2). Hence, these tryptic peptides are likely to be contaminants. Furthermore, the bimodal distribution of predicted binding affinities observed in HLA alleles whose motifs contain an arginine or a lysine at the last position, such as HLA-A*11:01 (Figure 2a), strongly suggests that some of the identified tryptic peptides are not true ligands for these alleles as well.

**Figure 2.**
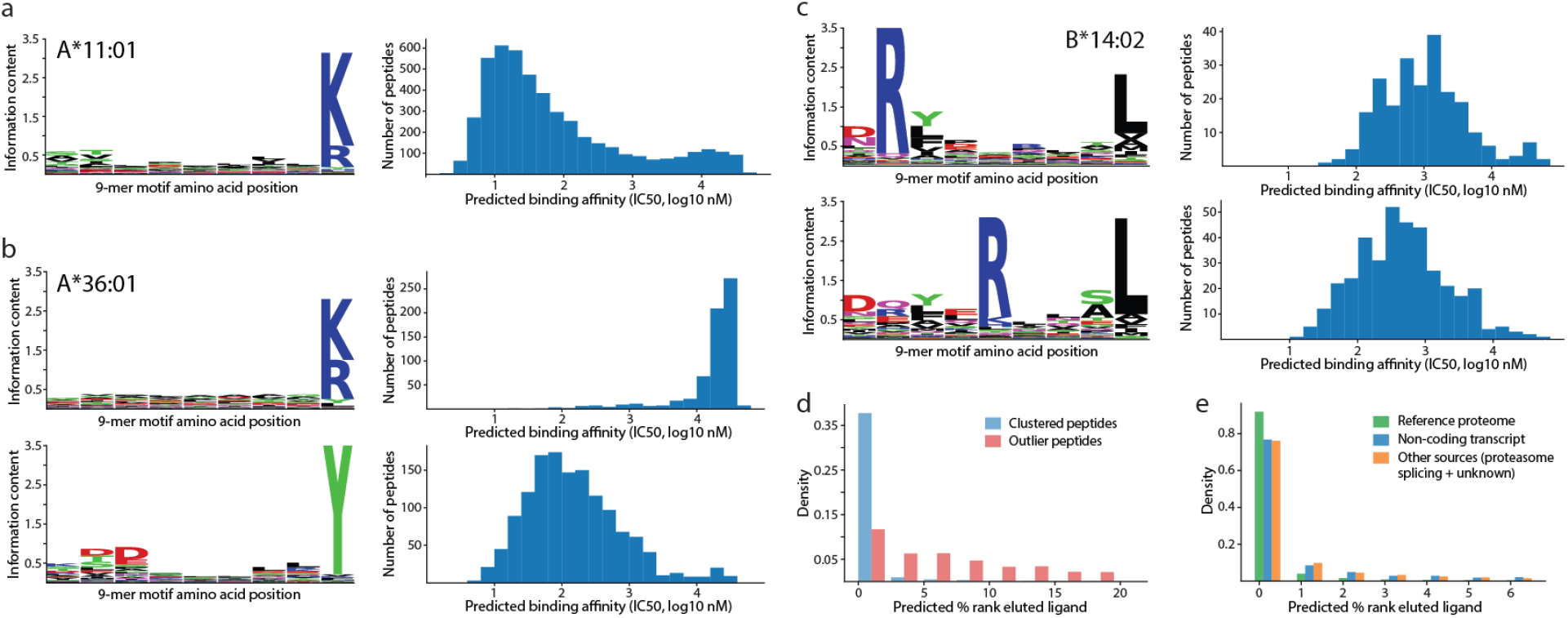
Unsupervised clustering revealed potential false positives and multiple motif specificities. a) Single 9-mer motif identified for HLA-A*11:01 together with predicted binding affinities (IC50, nM unit). b) Two motifs identified for HLA-A*36:01, one of which consists mainly of tryptic peptides and exhibits lower affinities (higher IC50 value indicates lower affinity). The top motif is expected to be a false positive. c) Two distinct motifs identified for HLA-B*14:02 with arginine at different residue positions but similar predicted affinities. d) Distributions of predicted percentage rank (% rank) of eluted ligand for clustered peptides and outlier peptides. A higher % rank indicates lower binding affinity. Bin size is 2%. e) Distributions of predicted percentage rank of eluted ligand for peptides from various sources. Bin size is 1%.

### HLA alleles with multiple binding motifs

In addition to revealing clusters of false-positive tryptic peptides, unsupervised motif clustering also showed that several HLA class I alleles possess multiple motif specificities that cannot be explained by length alone^11^. For example, antigens of HLA-B*14:02 contain arginine exclusively at either the 2^nd^ or the 5^th^ position of the motif with only slight differences in predicted binding affinities (Figure 2c, average predicted affinities are 2,067 nM and 1,733 nM, respectively). The motif for this allele was previously reported as a combined pattern with arginine at both positions^10,14^. Other alleles with multiple, clearly distinct motifs include HLA-B*15:01, HLA-B*51:01, and HLA-B*53:01 (Supplementary Figure 3). Additionally, several alleles also contain multiple related motifs that differ only by the shift of the anchor residue at the 2^nd^ position to the 1^st^ position (Supplementary Figure 4).

### False positives in HLA peptidomics data

A by-product of unsupervised motif clustering is the designation of outlier peptides that do not fit into any motif. Here, a peptide is labeled as an outlier if the quality of the motif clustering, as measured by Kullback-Liebler distance in GibbsCluster, is improved by removing the peptide from the analysis. This result revealed that up to 5-6% of identified peptides were classified as outliers for some HLA alleles (e.g., HLA-B*14:02 and HLA-A*02:05, Supplementary Table 2). As expected, the predicted binding percentage ranks of these outliers were much higher than those of peptides belonging to motif clusters (Figure 2d, higher percentage rank indicates weaker binding affinity). More than 83.8% and 95.5% of outliers do not pass the 2% rank threshold for weak binder and the 0.5% rank threshold for strong binder^24^, respectively. In contrast, only 10.2% and 20.4% of peptides that belong to motif clusters failed the same thresholds. Among peptides with unknown origins, which were identified solely by *de novo* sequencing, more than 47% of them pass the 0.5% rank threshold for strong binder (Figure 2e).

To test whether outlier peptides identified by unsupervised motif clustering are false positives or true ligands with very weak binding affinity, we performed an HLA binding assay on 59 newly identified antigens for HLA-B*14:02 (Supplementary Table 3, 13 outliers and 46 non-outlier peptides). This assay showed that all outlier peptides except LRNGGHFVI and LPFCRPGPEGQL exhibited almost no binding activity against the HLA molecules (Figure 3a, relative binding activity <1% of positive control). The high binding affinity of LRNGGHFVI and LPFCRPGPEGQL may be attributed to the arginine residues. LRNGGHFVI was likely called an outlier because its non-arginine residues did not fit the motif profile of HLA-B*14:02 (Figure 2c, top cluster). For LPFCRPGPEGQL, this peptide was likely called an outlier because the middle arginine residue was not predicted to take part in the 9-mer binding motif by NetMHCpan (the predicted core motif was LPFGPEGQL). Overall, the experimental binding result is in good agreement with computational affinity prediction (Figure 3b, Spearman’s rank correlation = –0.62 with p-value = 1.6e-7). These pieces of evidence together strongly suggest that outlier peptides are false positives.

**Figure 3.**
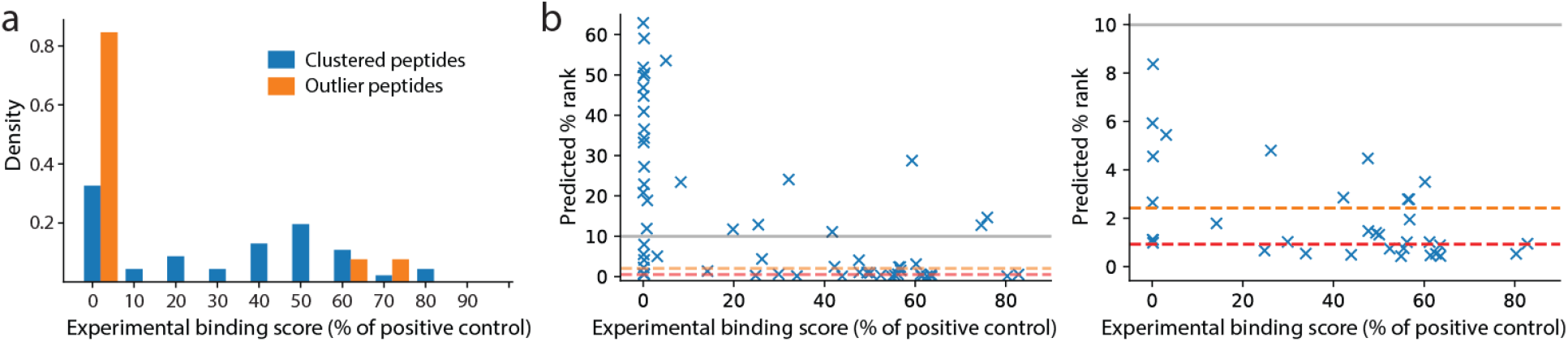
HLA binding assay for HLA-B*14:02. Peptide synthesis and binding assay were performed by ProImmune, Ltd. (see Methods). a) Distributions of binding scores, measured as the percentages of the binding activity compared to a positive control, for clustered peptides (n = 46) and outlier peptides (n = 13). b) Comparison of predicted percentage ranks of eluted ligand (% rank) and binding scores. The orange and red dashed lines indicate the 2% rank and 0.5% rank thresholds for weak and strong binders, respectively. The left panel shows the full range of % rank while the right panel shows the zoomed-in at % rank below 10%.

### Application of SMSNet on multi-allelic peptidomics data

To showcase the capability of SMSNet in a multi-allelic setting, SMSNet and PEAKS^25,26^ were used to analyze an HLA peptidomics experiment of a B-lymphoblastoid cell line expressing HLA-A*01:01, HLA-B*08:01, HLA-C*07:01, HLA-DPA1*01:03, HLA-DPB1*04:01/02:01, HLADQA1*05:01/05:01, HLA-DQB1*02:01/02:01, and HLADRB1*03:01/03:01. HLA class I and class II peptidomes were isolated and analyzed separately. NNAlign_MA^27^ was used to predict the binding probabilities for each identified antigen simultaneously against all HLA class I or class II alleles present. The maximum predicted binding score was taken for each peptide. Peptide sequencing with PEAKS was performed in two modes: the *de novo*-assisted database search mode (PEAKS-DB) and the fully *de novo* mode (PEAKS-DeNovo). As each tool was optimized differently, the confidence thresholds for peptide identification were set separately (see Methods). For PEAKS-DeNovo, confidence score thresholds ranging from 0.7 to 0.9 were explored. The results for PEAKS-DeNovo at a score threshold of 0.7 were selected, but it should be noted that increasing this threshold did not alter the conclusion.

For HLA class I peptidome, SMSNet and PEAKS-DB had a 40% overlap at peptide level (Figure 4a and Supplementary Table 4) and agreed on the same peptides for 98% of the MS/MS spectra identified by both tools (2,170 of 2,215 spectra). In contrast, PEAKS-DeNovo produced quite a different set of peptides (Figure 4a). SMSNet and PEAKS-DeNovo agreed on the same peptide for only 27% of the MS/MS spectra identified by both tools (526 of 1,973 spectra). To assess the quality of peptides identified by each tool, predicted HLA binding scores and peptide identification confidence scores were visualized together. Tools that identified peptides with high HLA binding scores with high confidences should be preferable. This analysis revealed that both SMSNet and PEAKS-DB identified peptides with high predicted binding probabilities and high confidences (heatmaps in Figure 4b). On the other hand, peptides identified *de novo* by PEAKS-DeNovo exhibited a bimodal distribution of predicted binding probabilities, with two modes at 0.5 and 1.0 (Figure 4c, the leftmost panels), which indicated that there is a substantial number of false positives.

**Figure 4.**
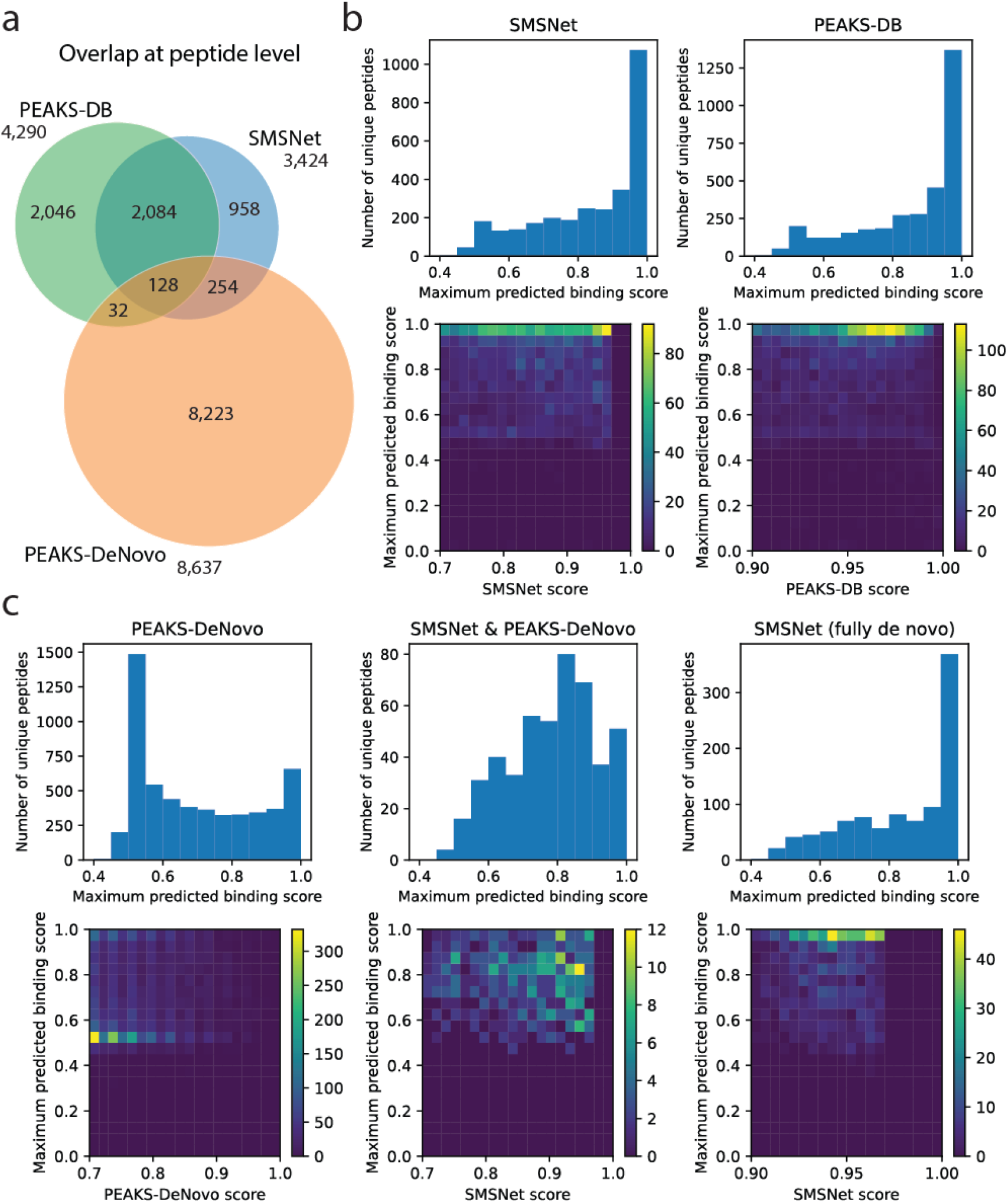
Comparison of SMSNet and PEAKS on multi-allelic HLA class I peptidomics sample. a) Overlap of identified peptides between SMSNet, the *de novo*-assisted database search mode of PEAKS (PEAKS-DB), and the fully *de novo* mode of PEAKS (PEAKS-DeNovo). b) Histograms show the distributions of predicted binding scores, calculated as the maximum score over HLA-A*01:01, HLA-B*08:01, and HLA-C*07:01 which are expressed in the cells, for peptides identified by SMSNet and PEAKS-DB. Heatmaps show the association between predicted binding scores and peptide identification confidence scores reported by each software. c) Similar visualizations for peptides identified by PEAKS-DeNovo, peptides identified in common by PEAKS-DeNovo and SMSNet, and peptides fully identified by the *de novo* sequencing step of SMSNet (SMSNet can identify the full sequences of some peptides without relying on reference database).

**Figure 4.**
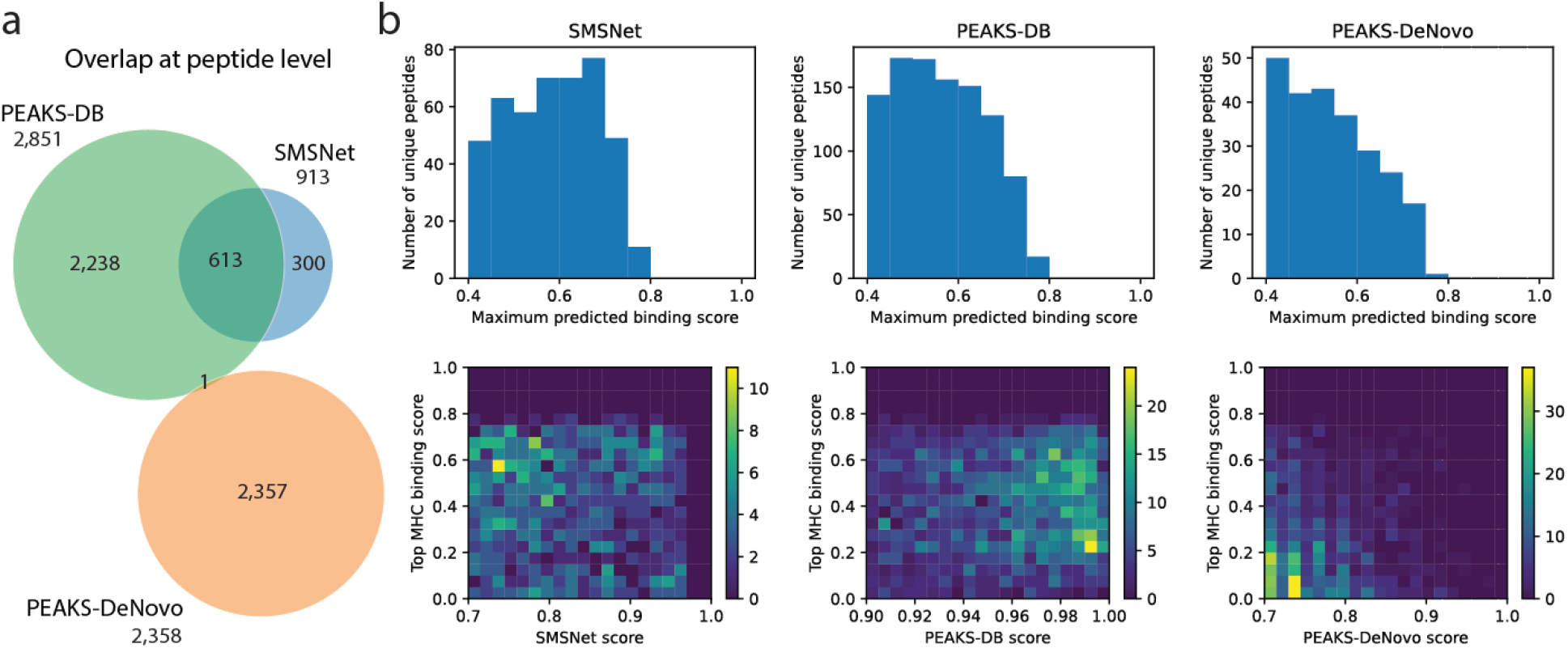
Comparison of SMSNet and PEAKS on multi-allelic HLA class II peptidomics sample. a) Overlap of identified peptides between SMSNet, the *de novo*-assisted database search mode of PEAKS (PEAKS-DB), and the fully *de novo* mode of PEAKS (PEAKS-DeNovo). b) Histograms show the distributions of predicted binding scores, calculated as the maximum score over HLA-DPA1*01:03, HLA-DPB1*04:01/02:01, HLADQA1*05:01/05:01, HLA-DQB1*02:01/02:01, and HLADRB1*03:01/03:01 which are expressed in the cells, for pepides identified by SMSNet, PEAKS-DB, and PEAKS-DeNovo. Heatmaps show the association between predicted binding scores and peptide identification confidence scores reported by each software.

To rule out the possibility that SMSNet produced peptides with high quality only because it relied on a follow-up database search after *de novo* sequencing to reduce errors, the set of peptides identified by both SMSNet and PEAKS-DeNovo and the set of peptides fully identified *de novo* by SMSNet before the database search step were analyzed separately. There were clear shifts in predicted binding scores toward 0.8-1.0 in both cases compared to PEAKS-DeNovo’s predictions (Figure 4c, the middle and rightmost panels), suggesting that *de novo* sequencing by SMSNet identified highly probable peptides. It should be noted that all methods also identified other peptides whose lengths do not match the expected lengths of HLA class I ligands (8-15 amino acids), and peptides with modifications were not considered here because their binding probabilities could not be predicted.

For the HLA class II peptidome, all tools made fewer identifications and had smaller overlap than HLA class I peptidome’s results (Figure 5a). This finding is likely because HLA class II antigens are much longer^28^ and consequently harder to confidently identify from MS/MS spectra. Only one peptide identified by PEAKS-DeNovo was also identified by others. In terms of the predicted binding scores, peptides identified by SMSNet exhibited slightly higher scores than PEAKS-DB’s (Figure 5b, Mann-Whitney p-value = 0.0131) and PEAKS-DeNovo’s (Mann-Whitney p-value = 4.34e-60). But as most of the predicted binding probabilities were below 0.5, it is inconclusive whether one tool is better than the others.

## Discussion

Our work highlighted the need for a careful downstream analysis of peptides identified from the HLA peptidomics experiment to remove potential false positives. Although a prior work has provided detailed analyses to account for non-ligand contaminants^20^, there are still true peptide identifications that bind very weakly or non-specifically to the target HLA allele. Inclusion of these peptides as true HLA ligands in community database can potentially mislead researchers as HLA peptidome-derived peptides are not accompanied with binding affinity values. Unsupervised clustering of identified putative HLA ligands not only elucidate allele-specific binding motif patterns^11,12^ but also revealed clusters of tryptic peptides for HLA alleles that should not recognize an arginine or a lysine at the C-terminus of the binding motif (Supplementary Figure 1) as well as outlier peptides that do not fit into any cluster. A small-scale HLA binding experiment of putative ligands of HLA*B14:02 confirmed that almost all outliers (11 of 13) exhibited no binding activity (Figure 3a, relative affinity < 1% of positive control) while 72% (33 of 46) of non-outliers exhibited some binding activities. Outlier peptides are also predicted to be weaker binders than *de novo*-identified peptides whose origins cannot be verified (Figure 2d and 2e, NetMHCpan % rank eluted ligand). Similarly, most tryptic peptides are likely false positives because their predicted binding affinities are not stronger than those between random tryptic peptides and HLA alleles (Supplementary Figure 2).

Overall, there are 3,846 potential false positives identified here that have been reported as positive antigens in the IEDB database. Although this number may seem small compared to the current size of the IEDB database (>300,000 allele-specific antigens), the presence of potential false positives is substantial for HLA alleles with fewer known ligands. For example, 23% (679 of 2,957), 16% (342 of 2,165), and 11% (209 of 1,843) of IEDB reported ligands for HLA-C*03:03, HLA-A*36:01, and HLA-B*57:01, respectively, are flagged as potential false positives here. Furthermore, the bimodal distribution of predicted affinities suggested that there are more false positives among peptides that belong to motif clusters (Figure 2a). Hence, careful analysis of both future HLA peptidomics data and the data already deposited into the IEDB database is needed in order to maintain the integrity of community antigen databases and prevent errors from propagating into HLA binding prediction and immunogenicity prediction models.

It is interesting to note that this work and prior unsupervised clustering analyses of the same HLA class I alleles^11,12^ do not always identify the same multiple motif specificities. For example, three motifs were identified for HLA-B*15:01 here (Supplementary Figure 3) but not in prior analysis^11^. On the other hand, three motifs for HLA-B*07:02 were previously reported^12^, but only a single motif was identified here. This latter case is especially unexpected because the motif identified here was not the one with the highest number of associated peptides among the three reported motifs. As a quality control, both motifs of HLA-B*51:01 (Supplementary Figure 3) were consistently identified^11^. In addition to multiple specificities, related motifs that differ by a shift of the 2^nd^ residue position to the 1^st^ residue position, with only minor changes in predicted binding affinities, were observed in several alleles (Supplementary Figure 4). These likely indicate the presence of 10-mer or longer motif patterns that were truncated to 9-mer during the core binding motif prediction by NetMHCpan. Lastly, unsupervised clustering was also able to capture minor inter-residue cooperation between non-anchor positions and represent them in separate motif clusters (HLA-B*53:01 in Supplementary Figure 3, HLA-B*15:03 and HLA-B*40:01 in Supplementary Figure 4).

Our work also illustrated the capability of hybrid *de novo* sequencing with SMSNet for uncovering new HLA antigens in both mono-allelic and multi-allelic peptidomics samples. More than 36,000 new peptide-HLA pairs were identified from public mono-allelic HLA class I peptidomics datasets^8,17^ that have already been extensively analyzed. The new putative antigens could potentially expand the antigen pools for some HLA alleles by up to 40% (Figure 1a and 1c). SMSNet exhibited good agreement with the *de novo*-assisted database search mode of PEAKS (PEAKS-DB), both producing peptide identifications with high predicted binding affinities to HLA class I alleles (Figure 4b). Furthermore, in the absence of a reference proteome database, SMSNet was able to produce peptides with higher predicted binding affinities than the *de novo* mode of PEAKS (PEAKS-DeNovo, Figure 4c). Putative HLA class II antigens identified by SMSNet also have slightly higher predicted binding affinities than both modes of PEAKS (Figure 5b), while PEAKS-DB produced many more identifications. The drop in the number of peptides identified from HLA class I to HLA class II peptidomics data is likely because HLA class II antigens consist of longer peptides which are more difficult to identify, especially for SMSNet and PEAKS-DeNovo which rely primarily on *de novo* sequencing. Overall, *de novo* analysis of HLA peptidomics would benefit from combining results from SMSNet and PEAKS-DB together to increase antigen detection sensitivity. It should be noted that combining results from multiple software tools is a well-established approach that has been shown to improve the quality of proteomics analyses^29,30^.

## Methods

### Cell line and antibody preparation

B-lymphoblastoid cell line (BLCL1408-1038) expressing HLA-A*01:01, HLA-B*08:01, HLA-C*07:01, HLA-DPA1*01:03, HLA-DPB1*04:01/02:01, HLADQA1*05:01/05:01, HLA-DQB1*02:01/02:01, and HLADRB1*03:01/03:01 was purchased from Fred Hutchinson Cancer Research Center, Washington, USA. Cells were cultured in RPMI 1640 media supplemented with 10% fetal bovine serum, 50 U/ml penicillin in a humidified incubator at 37C with 5% CO_2_. Purified pan HLA-A, -B, -C and pan HLA-DR, -DP, -DQ antibodies were generated from W6/32 (ATCC, USA) and IVA12 (provided by the lab of Professor Anthony Purcell, Monash University, Australia) hybridoma cells cultured in RPMI 1640 media supplemented with 10% fetal bovine serum, 50 U/ml penicillin and expanded in roller bottles at 37C with 5% CO_2_. Secreted monoclonal antibodies were harvested from spent media and purified using Protein A resin with ÄKTA purification system (Cytiva, USA).

### Immunoprecipitation of HLA class I and class II complexes

BLCL1408-1038 cell pellets (1 × 10^8^) were pulverised using an MM400 Retsch Mixer Mill (Retsch, Germany) and lysed with 0.1% IGEPAL CA-630, 100 mM Tris, 300 mM NaCl, pH 8.0 Complete Protease Inhibitor Cocktail (Roche, Switzerland). The supernatant was passed through a Protein G resin pre-column (500 μL) to remove non-specific binding materials. HLA class I and II immunoaffinity purification was performed as previously described^31^. Briefly, the pre-cleared supernatant was incubated with 10 mg of pan HLA-A, -B, and -C antibodies or 10 mg of pan HLA-DR, -DP, and -DQ antibodies coupled to Protein G resin with rotation overnight at 4C. After conjugation, the resins were washed with 10 ml of ice-cold wash buffer 1 (0.005% IGEPAL, 50 mM Tris, pH 8.0, 150 mM NaCl, 5 mM EDTA), 10 ml of ice-cold wash buffer 2 (50 mM Tris, pH 8.0, 150 mM NaCl), and 10 ml of ice-cold wash buffer 3 (50 mM Tris, pH 8.0, 450 mM NaCl). Bound complexes were eluted from the column using 5 column volumes of 10% acetic acid. Eluted peptides were fractionated by reverse-phase high-performance liquid chromatography (Shimadzu, Japan) on a 4.6 mm diameter Chromolith SpeedROD RP-18 (Merck, USA). The optimized conditions were as follows: mobile phase A (0.05% v/v TFA, 2.5% v/v ACN in water), mobile phase B (0.045% v/v TFA, 90% v/v ACN in water), flow rate of 1 mL/minute, temperature of 30C, and injection volume of 200 μL. The elution program was set as follows: 0-5% of mobile phase B over 1 minute, 5-15% of mobile phase B over 4 minutes, 15-45% of mobile phase B over 30 minutes, 45-100% of mobile phase B over 15 minutes, and 100% of mobile phase B over 4 minutes. Fractions were collected in 1 mL each. Consecutive fractions were pooled into 11 fractions. Pooled fractions were concentrated by vacuum centrifugation and reconstituted in 0.1% FA.

### LC-MS/MS analysis of HLA peptidome

Pooled peptide fractions eluted from an HLA class I sample and an HLA class II sample were analyzed on a Q Exactive mass spectrometer (Thermo Fisher Scientific, USA) coupled to an EASY-nLC 1000 (Thermo Fisher Scientific, USA). Peptide samples were separated at a flow rate of 300 μL/minute of buffer B (80% ACN, 0.1% FA). The gradient was set at 4-20% of buffer B over 30 minutes, 20-28% of buffer B over 40 minutes, 28-40% of buffer B over 5 minutes, 40-95% of buffer B over 3 minutes, washing with 95% of buffer B over 8 minutes, re-equilibration with buffer A (2% ACN/0.1% FA) over 5 minutes. Mass spectra resolutions were set at 70,000 for full MS scans and 17,500 for MS/MS scans. The normalized collision energy for HCD fragmentation was set at 30%. The m/z scan range was set at 350-1,400. Dynamic exclusion was set at 15 seconds. For HLA class I samples, the maximum injection times were set at 120 ms for full MS scan and 120 ms for MS/MS scans. Precursor ions with charge states +2, +3, +4, and +5 were accepted. For HLA class II samples, the maximum injection times were set at 200 ms for full MS scan and 120 ms for MS/MS scans. Precursor ions with charge states +2, +3, +4, +5, and +6 were accepted.

### Collection of published HLA class I peptidomics and antigen data

A combined dataset of mass spectrometry raw data of mono-allelic HLA class I peptidomes (399 raw files, 88 HLA alleles) were obtained from two prior studies^8,17^ (MSV000080527 and MSV000084172). List of reported antigen-HLA pairs were obtained from the Immune Epitope Database^15^ (IEDB, downloaded December 2020), the HLA Ligand Atlas^32^ (downloaded June 2020), and from peptidomics analyses of multi-allelic patient samples^8,33^. It should be noted that these recent studies of patient samples not only reported new data but also provided compilations of multi-allelic peptidomics data from earlier studies.

### Peptide sequencing of MS/MS data

For *de novo* peptide sequencing with SMSNet^21^, MS/MS spectra and precursor masses were extracted from raw MS files using ProteoWizard^34^ with the following parameters: Peak Picking = Vendor for MS1 and MS2, Zero Samples = Remove for MS2, MS Level = 2-2, and the default Title Maker. Charge state deconvolution was not performed. The SMSNet-M model which treats carbamidomethylation of cysteine as fixed modification and oxidation of methionine as variable modification was used. Target amino acid-level false discovery rate was set at 5%. Precursor mass tolerance of 30 ppm was applied to discard identified peptides with high mass deviations. Partially identified peptides were searched against a UniProt^35^ reference human proteome (downloaded August 2020) and a GRCh38 RefSeq^36^ non-coding transcriptome (downloaded August 2020) to fill in the missing amino acids. From the transcriptome data, possible open reading frames that translate to at least 5 amino acids in length were considered.

For database search and *de novo* peptide sequencing with PEAKS version 8.5^25^, raw MS files were searched against a UniProt reference human proteome and reversed decoys. Cleavage enzyme specificity was set to none. Carbamidomethylation of cysteine, oxidation of methionine, and phosphorylation of serine, threonine, and tyrosine were set as variable modifications. A maximum of three modifications per peptide were allowed. Mass tolerances were set at 10 ppm for precursor mass and at 0.02 Da for fragment mass. Target peptide-level false discovery rate was set at 1%.

### Explaining peptides with unknown origins

Peptides that do not match to either reference human proteome or non-coding transcriptome were further analyzed to explain their origins. The proteasome-mediated splicing mechanism, which causes the joining of two distal peptide fragments from the same protein into a new contiguous peptide, was explored by considering all possible combinations of 3-12 amino acid peptides originating from non-overlapping regions of each protein. Only proteins that were already identified with some peptides in the same dataset were considered as sources of spliced peptides. If multiple possible splicing events could explain an observed peptide, the one involving peptides that are nearest to each other on a protein was selected as the most likely explanation. To check whether some peptides could be explained by splicing events by chance alone, the amino acid sequences of these peptides were randomly shuffled and reanalyzed. This revealed that using proteasome-mediated splicing as explanation may not be reliable because as many as 45% (99 out of 222) of randomized sequences could still be matched to some hypothetical spliced peptides. Furthermore, missense mutations could serve as an alternative explanation for 30% (66 out of 222) of peptides that could be explained by proteasome-mediated splicing.

### HLA binding affinity and binding motif analyses

For peptides identified from mono-allelic HLA peptidome experiments^8,17^, the binding affinities and the 9-mer binding motifs for the corresponding HLA alleles were predicted using NetMHCpan-4.1^24^ with default setting. For peptides identified from multi-allelic B-lymphoblastoid cell line, the binding affinities were predicted against all HLA class I or class II alleles present using NNAlign_MA^27^. Predicted 9-mer binding motifs for each HLA class I allele were then clustered using GibbClusters^23^. For each allele, the clustering was performed with number of clusters ranging from 1 to 5, with or without outlier detection, and with inter-cluster penalty parameter λ ranging from 0.1 to 0.8. The optimal number of clusters was determined from the parameter setting with the highest Kullback-Liebler distance (KLD) as recommended by the authors^23^. Information contents and the amino acid profiles of 9-mer binding motif clusters were visualized using Logomaker^37^.

### HLA binding assay

The binding activities of selected 59 newly identified candidate antigens for HLA-B*14:02 (Supplementary Table 3) were assessed using the REVEAL MHC-peptide binding assay provided by ProImmune, Ltd. (Oxford, UK). Peptides were synthesized and quality checked using MALDI-TOF mass spectrometry by ProImmune, Ltd. (Oxford, UK). Binding activities were reported as percentage relative to the affinity of a positive control (a known high-affinity T cell epitope for HLA-B*14:02). According to the experiment report provided by the company, the standard error of the reported affinities is 3 percentage points.

## Data availability

Identified peptides from public mono-allelic HLA peptidomes are provided in Supplementary Table 1 along with binding affinity prediction and outlier detection result. HLA binding assay results are provided in Supplementary Table 3. Identified peptides from the multi-allelic B cell peptidome are provided in Supplementary Table 4. Raw mass spectrometry data for the multi-allelic B cell peptidome are available at PXD028088. Visualizations of all identified motifs are available on FigShare at 10.6084/m9.figshare.16025226.

## Acknowledgements

This work was supported by the Thailand Research Fund MRG6280189 (S.S.), the Grant for Special Task Force for Activating Research, Ratchadapisek Sompoch Endowment Fund, Chulalongkorn University (S.S.), the Grant for the Development of New Faculty Staff, Ratchadapisek Sompoch Endowment Fund, Chulalongkorn University (S.S.), the Thailand Research Fund for Career Development Grant RSA6280026 (T.P.), and Program Management Unit for Competitiveness Grant C10F630106 (T.P.). We would like to thank Prof. Vorasuk Shotelersuk, Department of Pedratrics, Faculty of Medicine, Chulalongkorn University, for mentorship under the Thailand Research Fund program and Dr. Pokrath Hansasuta, Department of Microbiology, Faculty of Medicine, Chulalongkorn University for insightful advices on HLA research.

## Contributions

C.S., T.B., and P.S. analyzed HLA peptidomics data. P.M. performed experiments. S.S., P.S., and C.S. wrote the manuscript draft. S.S. and T.P. conceived and supervised the research. All authors contributed to and approved of the final manuscript.

## Competing interests

The authors declare no competing interest.

## Tables

None

## Supplementary Tables

Supplementary Table 1 – List of all identified peptides together with predicted binding affinities and outlier detection results from mono-allelic peptidomics data of 88 HLA class I alleles

Supplementary Table 2 – Percentages of outlier peptides for HLA class I alleles with low percentage of tryptic peptides

Supplementary Table 3 – HLA-B*14:02 binding assay results for selected 59 peptides

Supplementary Table 4 – SMSNet and PEAKS identification results for multi-allelic B-lymphoblastoid cell line

## Supplementary Figures

**Supplementary Figure 1.**
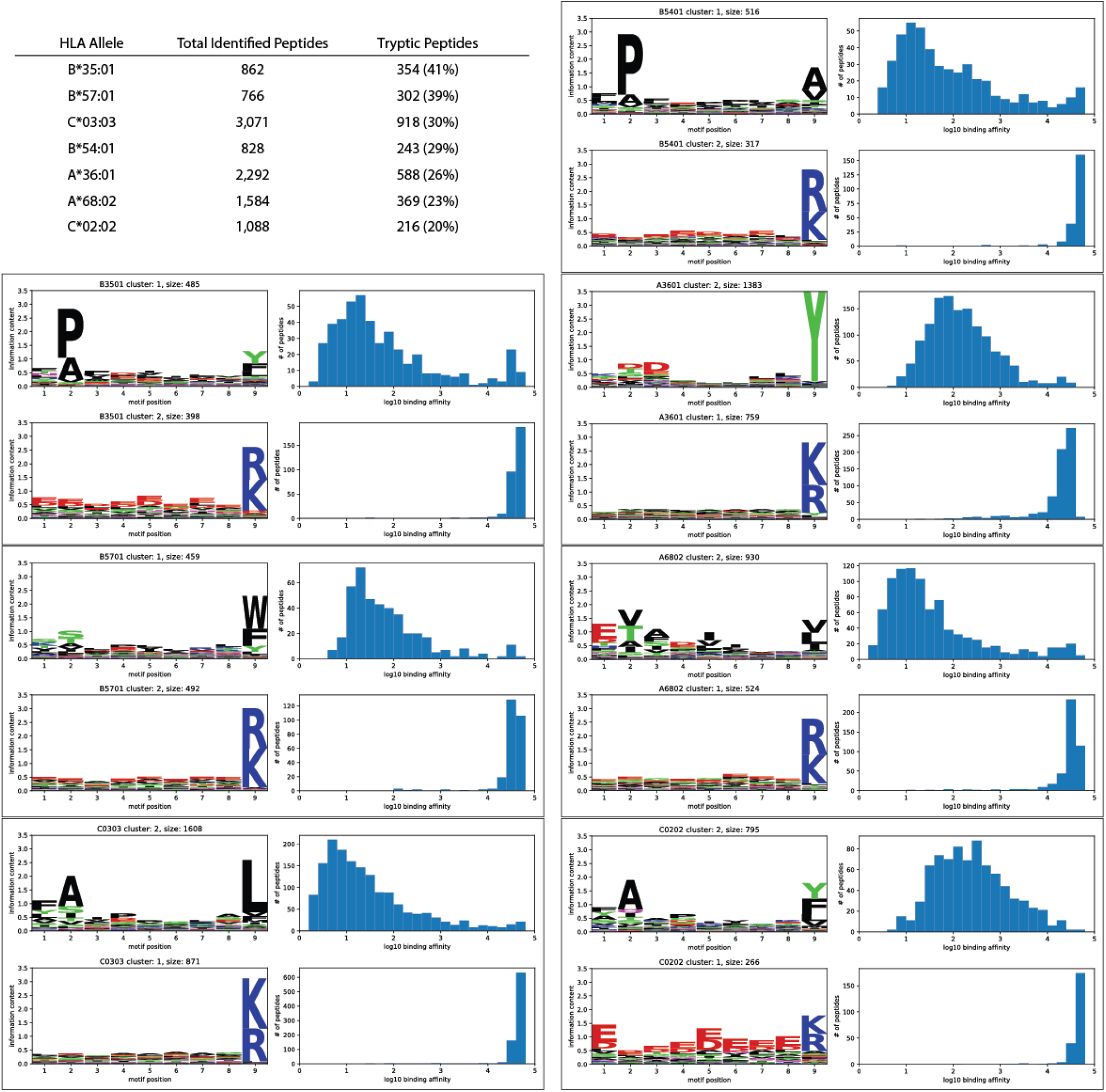
Extents of tryptic peptide contaminations in HLA peptidomics data. Data for the top 7 alleles with more than 20% contaminations are shown. The table lists the numbers of all identified peptides and tryptic peptides for each allele. Each boxed region contains the 9-mer motif profiles and distributions of predicted binding affinity for each allele, sorted in the same order as shown in the table from left to right.

**Supplementary Figure 2.**
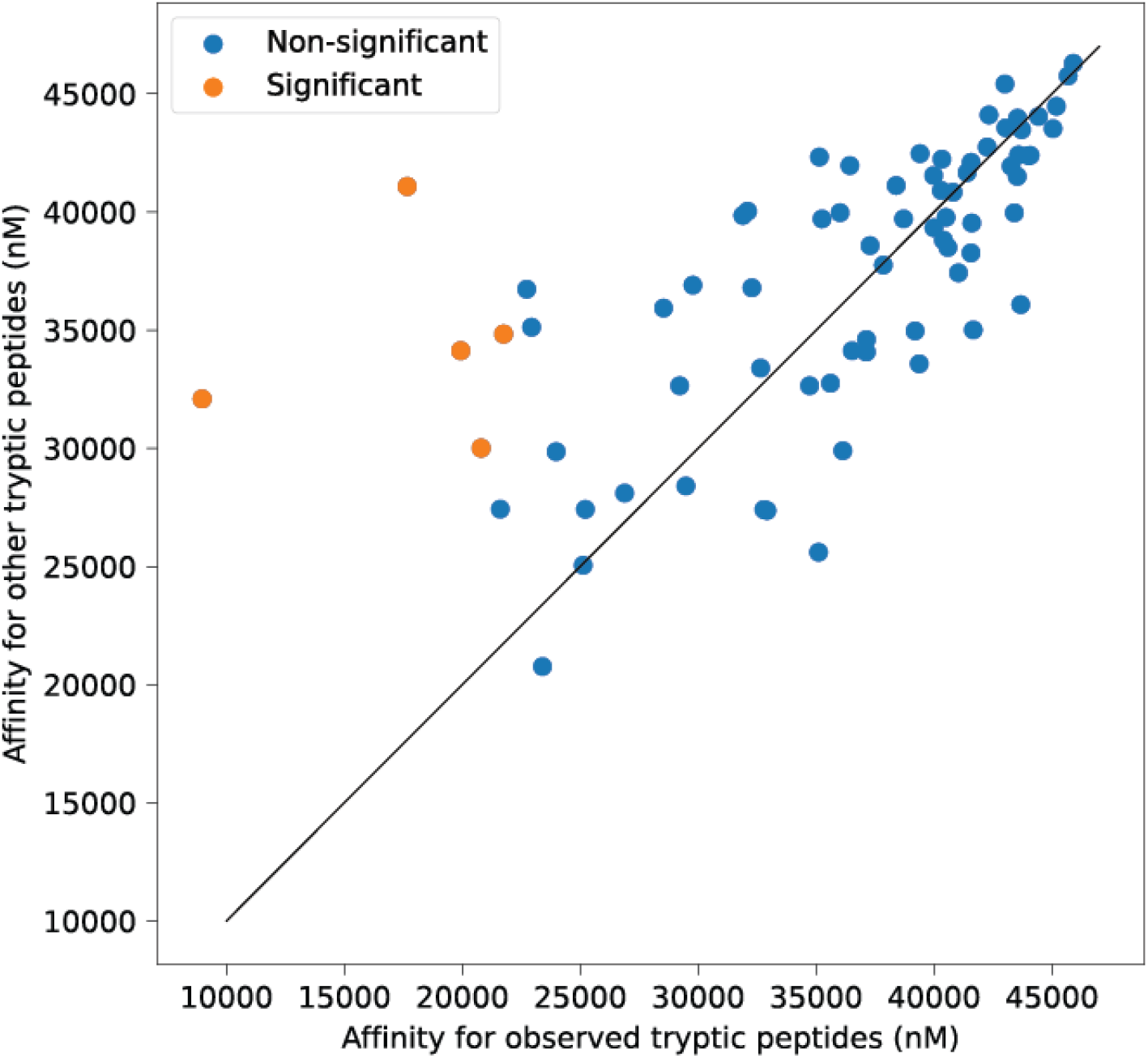
HLA alleles do not exhibit stronger affinities toward observed tryptic peptides than toward random tryptic peptides. Scatter plot shows the median predicted binding affinity (IC50, nM unit) between observed tryptic peptide-HLA allele pairs (x-axis) and that between random tryptic peptide-HLA pairs. Each data point represents one HLA allele. Higher IC50 value indicates lower affinity. Random tryptic peptides were selected from observed tryptic peptides in peptidomics data of all HLA alleles. Orange data points indicate the few HLA alleles that exhibit significantly stronger affinities toward tryptic peptides identified from the corresponding peptidomics data (Benjamini-Hochberg adjusted Mann-Whitney U test p-value < 0.05).

**Supplementary Figure 3.**
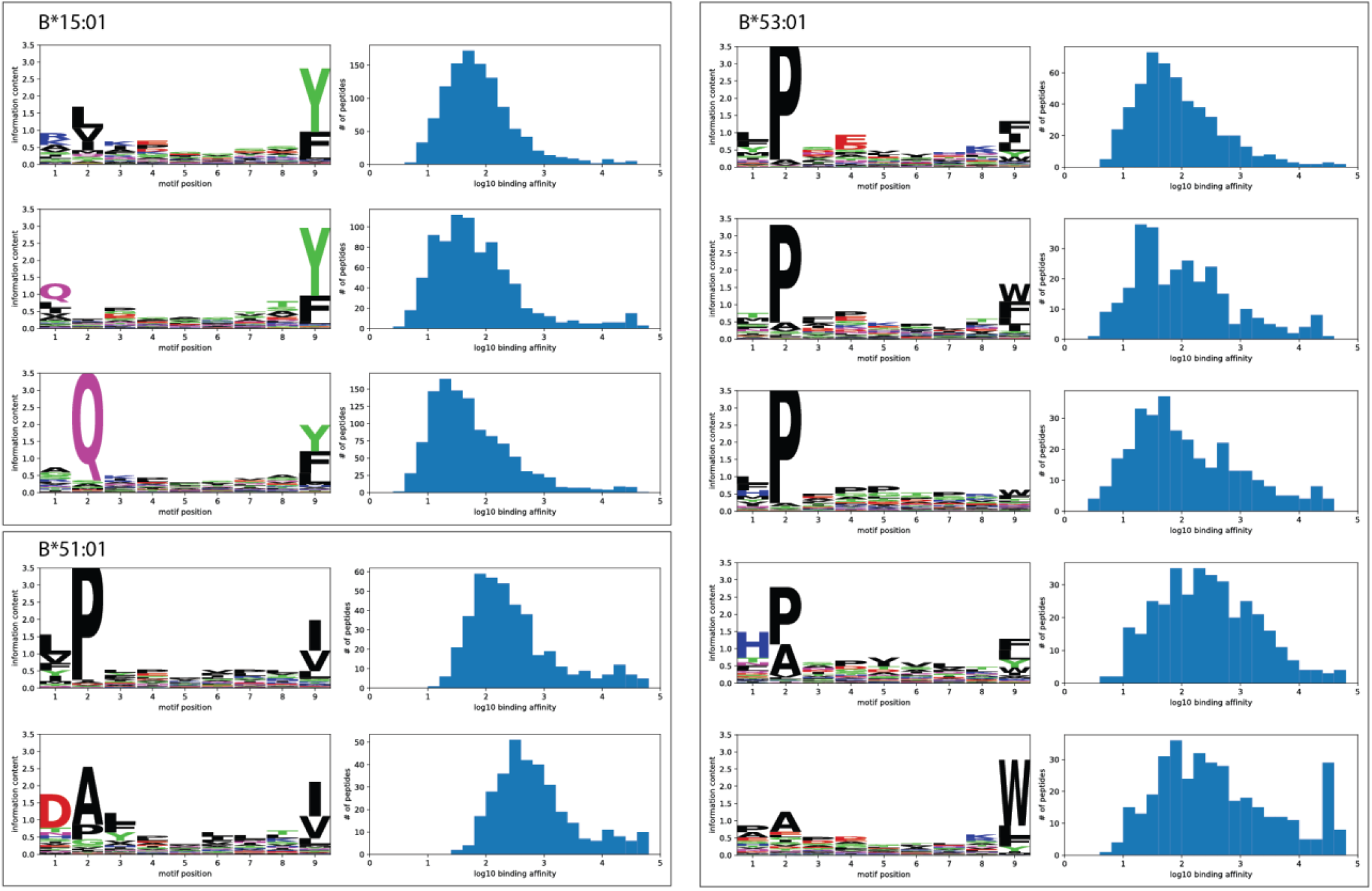
HLA alleles with multiple, clearly distinct motif specificities. Each boxed region contains motifs of the indicated HLA allele. Each 9-mer motif is shown alongside the distribution of predicted binding affinity (IC50, nM unit).

**Supplementary Figure 4.**
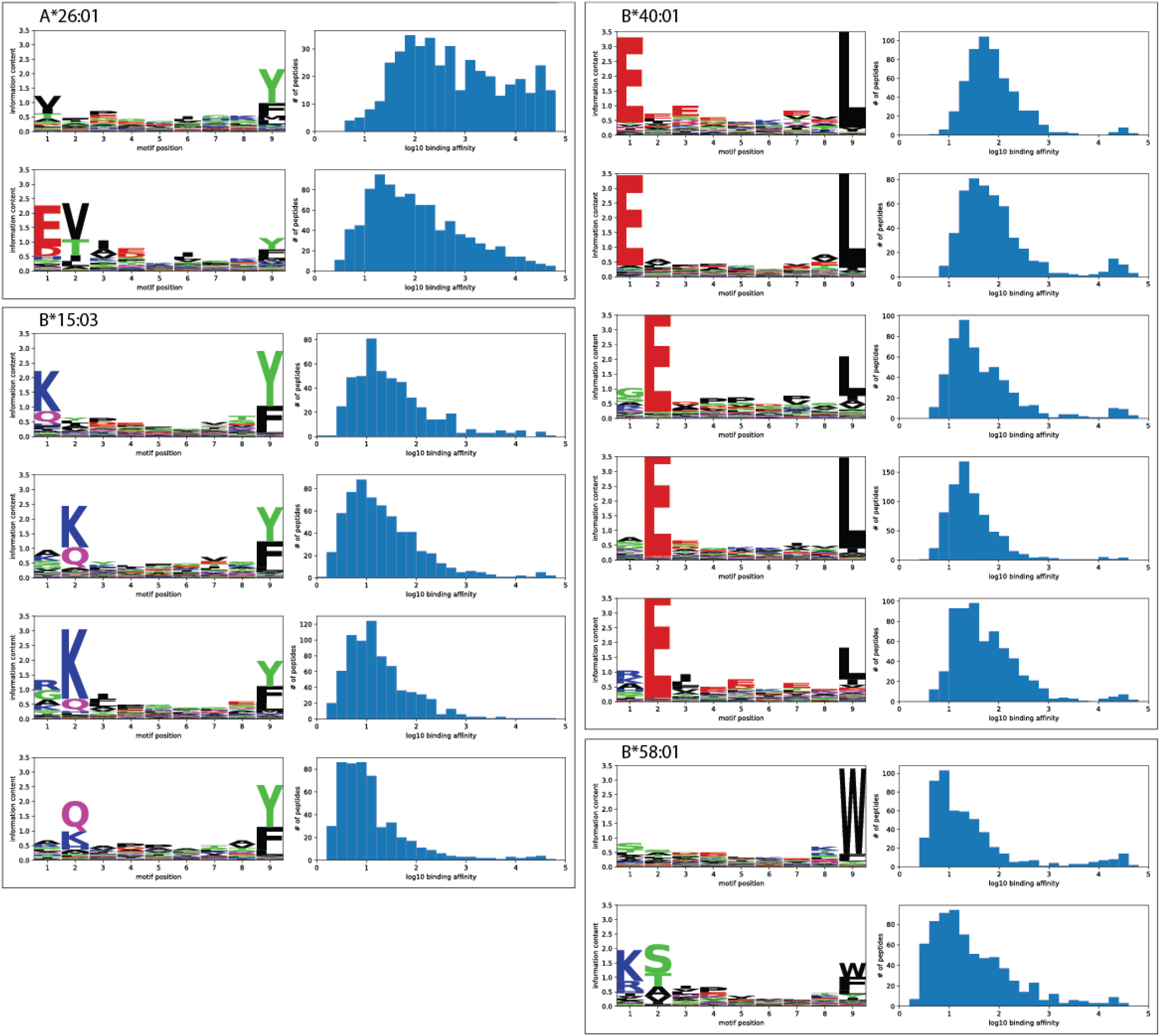
HLA alleles with multiple related motif specificities. Each boxed region contains motifs of the indicated HLA allele. Each 9-mer motif is shown alongside the distribution of predicted binding affinity (IC50, nM unit). These motifs possess similar anchor residues at the 2^nd^ position or shifted to the 1^st^ position.

## Notes

### Competing Interest Statement

The authors have declared no competing interest.

